# Dynamic Consequences of Specificity within the Cytidine Repressor DNA-Binding Domain

**DOI:** 10.1101/2021.02.28.433298

**Authors:** Colleen L. Moody, Jenaro Soto, Vira Tretyachenko-Ladokhina, Donald F. Senear, Melanie J. Cocco

## Abstract

The *E. coli* cytidine repressor (CytR) is a member of the LacR family of bacterial repressors that regulates nine operons with distinct spacing and orientations of recognition sites. Understanding the structural features of the CytR DNA-binding domain (DBD) when bound to DNA is critical to understanding differential mechanisms of gene regulation. We previously reported the structure of the CytR DBD monomer bound specifically to half-site DNA and found that the DBD exists as a three-helix bundle containing a canonical helix-turn-helix motif, similar to other proteins that interact with DNA [Moody, et al (2011), Biochemistry **50**:6622-32]. We also studied the free state of the monomer and found that since NMR spectra show it populates up to four distinct conformations, the free state exists as an intrinsically disordered protein (IDP). Here, we present further analysis of the DBD structure and dynamics in the context of full-site operator or nonspecific DNA. DBDs bound to full-site DNA show one set of NMR signals, consistent with fast exchange between the two binding sites. When bound to full-length DNA, we observed only slight changes in structure compared to the monomer structure and no folding of the hinge helix. Notably, the CytR DBD behaves quite differently when bound to nonspecific DNA compared to LacR. A dearth of NOEs and complete lack of protection from hydrogen exchange are consistent with the protein populating a flexible, molten state when associated with DNA nonspecifically, similar to fuzzy complexes. The CytR DBD structure is significantly more stable when bound specifically to the *udp* half-site substrate. For CytR, the transition from nonspecific association to specific recognition results in substantial changes in protein mobility that are coupled to structural rearrangements. These effects are more pronounced in the CytR DBD compared to other LacR family members.

## INTRODUCTION

The cytidine repressor protein (CytR), a member of the LacR family of bacterial repressors [1] [2] regulates multiple operons whose genes encode transport proteins and enzymes involved in ribonucleotide salvage and catabolism. Recently, CytR has been shown to control the production of flagella in uropathogenic *E. coli* that cause urinary tract infections. In *V. cholerae*, CytR is involved in nutrient sensing and controls competence [3-6], adhesion [7] and biofilm production [8].

In contrast to other transcription regulators, CytR tolerates widely variable spacing between the inverted repeats that comprise its natural operators, ranging from zero base pairs at *deoP2* to eleven base pairs at *rpoH*, a variation that accounts for up to a 38Å translation and a complete rotation of the half-sites. We have shown that this accommodation requires distinctly different thermodynamic modes of CytR-DNA binding, which we interpret as ranging from a relatively rigid conformation of CytR bound to zero base pair spacing at *deoP2* to a flexible and dynamic conformation bound to the nine base pair spacing of *nupG* [9, 10]. The *udp* promoter is involved in the regulation of expression of uridine phosphorylase [11] and is an example of an intermediate spacing of three base pairs between CytR recognition sites.

To address the structural mechanism underlying these effects we have used NMR spectroscopy to investigate the structure and dynamics of isolated DBD and linker peptide bound to operator DNA and when associated with nonspecific DNA. Previously we determined the structure of the CytR DBD monomer bound specifically to one DNA half-site of the uridine phosphorylase (*udp*) operator [12]. The CytR DBD structure demonstrated that the conformation of specifically-bound protein is different from the common conformation of the other four LacR DBDs whose structures are known. If helix 2 is superimposed, both helices 1 and 3 move in the CytR DBD compared to LacR.

We also showed that operator-DNA binding is accompanied by a substantial disorder to order transition in going from the free state to the three-helix bundle that comprises the DBD core. In contrast, the three helices of the LacR DBD remain largely folded regardless of whether this domain is free in solution or bound to DNA. We found that the free CytR DBD is intrinsically disordered, existing in multiple conformations with only 20% helicity as measured by circular dichroism and very few NOEs. We found two sets of NMR signals in helices 1 and 2 and four sets of signals in helix 3 (figure 6 in [12]). For a residue in multiple conformations, the NMR timescale of exchange is defined by the separation between NMR resonance peaks in cycles/sec (Hz). Measuring the distance in Hz between peaks sets an upper limit for the rate of exchange. We calculate that, based on the largest peak separation, the rate of exchange between species must be slower than 0.044 sec^−1^ when the DBD is free in solution. Thus, the alternate conformations of the DBD are stable and could represent unique conformations optimized for different operator structures. Thus, the CytR DBD can be characterized as an intrinsically disordered protein (IDP). Since our CytR DBD structure was published, it has been used as a paradigm in understanding IDPs in comparison to other folded LacR DBDs [7, 13-18].

Here we find that although isolated DBD also binds nonspecific DNA with only moderately lower affinity, this binding is not accompanied by the same magnitude of folding of binding to specific site DNA. In this respect, the CytR DBD with nonspecific DNA is a fuzzy complex where the protein is only partially folded [19-21]. Further, even when DBD is bound to full-site operator-DNA, the CytR hinge helix does not fold and a dimer interface is not formed as it is in LacR. We thus conclude that specific DBD-DNA contacts are necessary for full stabilization of the bound structure with folding of the DBD being coupled to DNA binding.

## MATERIALS AND METHODS

### Protein expression and purification

The 67 residue CytR DNA binding domain (DBD) was expressed in *E. coli* strain BL21(DE3)pLysS transformed with expression plasmid pSS584DBD[22]. ^15^N labeled protein was produced by growing the cells in Neidhardt’s minimal media [23] containing ^15^N-ammonium chloride as the sole nitrogen source and purified as described [24]. Purity was above 95% as assessed by SDS PAGE and MALDI mass spectrometry. Concentration was determined using extinction coefficient A_280_ = 0.202 (g/L)^−1^ cm^−1^ calculated from the amino acid composition [25].

### Oligonucleotide preparation

Deoxyribonucleotides were purchased as HPLC purified products from IDT Technologies, San Diego, CA.. Complimentary single strand oligonucleotides were annealed to double strands as described previously [24]. Each double strand sequence contained one strand conjugated to either Alexa 532 or Oregon Green 514 fluorescent dye through a 6-carbon linker to the 5’-phosphate.

### DBD binding to DNA by steady-state fluorescence anisotropy

Steady-state fluorescence measurements were made using an SLM 8100 equipped with a double-grating excitation monochromator and a single-grating emission monochromator. Fluorescence was excited at 530 nm and emission measured at 580 nm, both with excitation 4 nm and emission 8 nm band pass. Fluorescence intensity was recorded for 2 minutes at each position of the excitation and the emission polarizers. Anisotropy was calculated from the average intensities according to

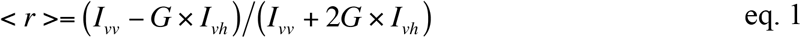

*I*_*vv*_ and *I*_*vh*_ are the intensities at the excitation polarizer in the vertical position and emission in the vertical and horizontal positions of the emission polarizer, respectively. The *G* factor,

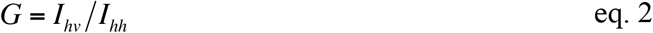

was calculated from the average of several measurements of free labeled DNA.

Binding titrations were conducted in 50 mM phosphate buffer, pH 6 containing 30 mM NaCl and 1 mM EDTA. Aliquots of DBD from a 5 mM concentrated stock were added to a solution of Alexa 532-labeled DNA (∼100 nM) in 2×10 mm quartz cuvettes at a temperature of 20±0.1 °C maintained by a Peltier device. After each addition of protein, the cuvette was incubated in the thermo-regulated cell holder for 10 minutes prior to taking intensity measurements to ensure that equilibrium had been attained. Anisotropy values were analyzed as a function of DBD concentration using non-linear least squares analysis in Origin 7 according to:

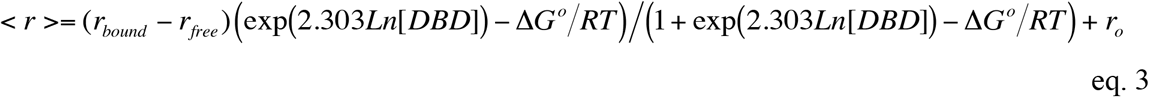

where *r*_*o*_ is a free labeled DNA anisotropy in the absence of protein, *r*_max_is the anisotropy of DBD-DNA complex, Δ*G*°is a standard free energy of binding, and *R* and *T* are the gas constant and absolute temperature. Dissociation constants are calculated as *k*_*d*_ = 1/*K*_*a*_ = exp(Δ*G*°*RT*).

#### Chemical shift analysis

Calculation of chemical shift changes, Δδ_amide_ was calculated using the changes in chemical shift of ^1^H and ^15^N according to the following equation from Williamson, 2013 [26]:

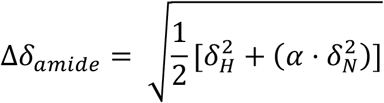

Where δ_*H*_ is the change in chemical shift of ^1^H and δ_*N*_ is the change in chemical shift of ^15^N between CytR DBD in the presences of specific and non-specific DNA fragments. α is the nitrogen scaling factor of 0.14 [26].

### NMR dynamics measurements

All NMR experiments were performed with samples in 90%H2O/10%D2O in 50 mM sodium phosphate, 30 mM NaCl, and 1 mM EDTA at pH 6. Samples with *udp* full-site contained 0.411 mM ^15^N CytR DBD and 0.889 mM natural abundance DNA. Samples with nonspecific DNA contained 0.4 mM ^15^N CytR DBD and 0.6 mM natural abundance half-site nonspecific DNA. The free DBD samples contained 1 mM DBD ^15^N CytR DBD.

NMR data were acquired on a Varian Inova 800MHz spectrometer. Data processing was performed using NMR Pipe (30) and spectra were analyzed using CCPNMR Analysis (31). Assignments were previously published[24]. ^15^N T1 and T2 relaxation measurements were made using ^15^N HSQC pulse sequences and varying relaxation times (46, 47) (T1 = 0, 50, 100, 200, 750, and 1500 ms; T2 = 0, 10, 30, 50, 70, 90, 110, 130, 150, 170, 190, 210, 230, and 250 ms). Peak intensities were fit to single exponential functions to determine the relaxation times (errors were calculated from the fit). Steady-state NOE values were calculated as the ratio between the peak intensities from the saturated and unsaturated spectra (48).

Hydrogen exchange measurements were performed by lyophilizing ^15^N CytR DBD with natural abundance DNA in standard buffer and then resuspending the sample in D_2_O. ^1^H-1D and HSQC experiments were acquired immediately upon resuspension and transfer to Shigemi NMR tubes. ^1^H-1Ds were acquired in between each HSQC (nt=16, ni=40, np=1024) for calibration purposes. HSQCs were processed without apodization and peak intensities were fit to single exponential curves to extract hydrogen-deuterium exchange rates for backbone amides in the CytR DBD *udp* half-site sample. Backbone amide protons of CytR DBD with nonspecific DNA exchanged too rapidly to obtain quantitative rate information. Protection factors were calculated for the CytR DBD with *udp* half-site sample using the following equation: Protection Factor = k_intrinsic_/k_measured_. The intrinsic rates were calculated based on studies by Bai et al. [27].

## RESULTS

We characterized the structure and dynamics of the CytR DBD in the context of three types of DNA substrates. These are: bound in a 1:1 complex to an operator half-site (*udp*), bound to a nonspecific DNA sequence (the same size as the *udp* half-site), and bound to full-length operator (*udp*), with the possibility that the DBD might dimerize on the DNA. CytR operators are comprised of inverted repeats of an eight base pair recognition motif [28] separated by variable spacing ranging from zero to nine base pairs [10]. These experiments used the natural operator (CytO) from *udp* (5’-GTAAATTTATGCAAC<u>GCA</u>TTTGCGTCATGGTG-3’), an operator with three base pair spacing (underlined) hereafter referred to as full-site DNA. For the single recognition site sample, the operator left half-site of *udp* (5’-ATTTATGCAACGCA −3’) is the best match to the consensus recognition motif defined by SELEX [28]. The nonspecific DNA substrate (5’ - AGCATGCTGGGGAC - 3’) is the same length as the *udpP* half-site (14 bp) but contains the binding sequence of an unrelated transcription factor, i.e., the human p53 binding sequence found in the *gadd45* promoter [29]. As **Figure 1** indicates, the NMR spectrum of the DBD bound to the full-site operator has many similarities to that of the half-site. When chemical shift changes between half- and full-site DNA spectra are plotted onto the monomer structure diagram in **Figure 1**, it appears that the largest changes in chemical environment are proximal to the first helix. In the presence of the nonspecific DNA, the DBD gives one set of signals at low temperatures (20 °C), but similar to the free protein [24], is mostly unfolded at 35 °C (**Figure 2**). Although the DBD is thermodynamically less stable when bound to nonspecific DNA then when bound to *udpP*, binding still produces a single structure. However, a comparison of HSQC spectra in Figure 2 and Figure 1B of [24] reveals that the chemical shifts of the protein/nonspecific DNA complex are different when compared to those of the protein/*udp* half-site (calculated in **Figure 3**). Comparison of spectra presented here and previously [24], shows that many DBD NMR peak positions are unique to the particular state of the protein (free, bound to specific DNA, or bound to nonspecific DNA).

**Figure 1.**
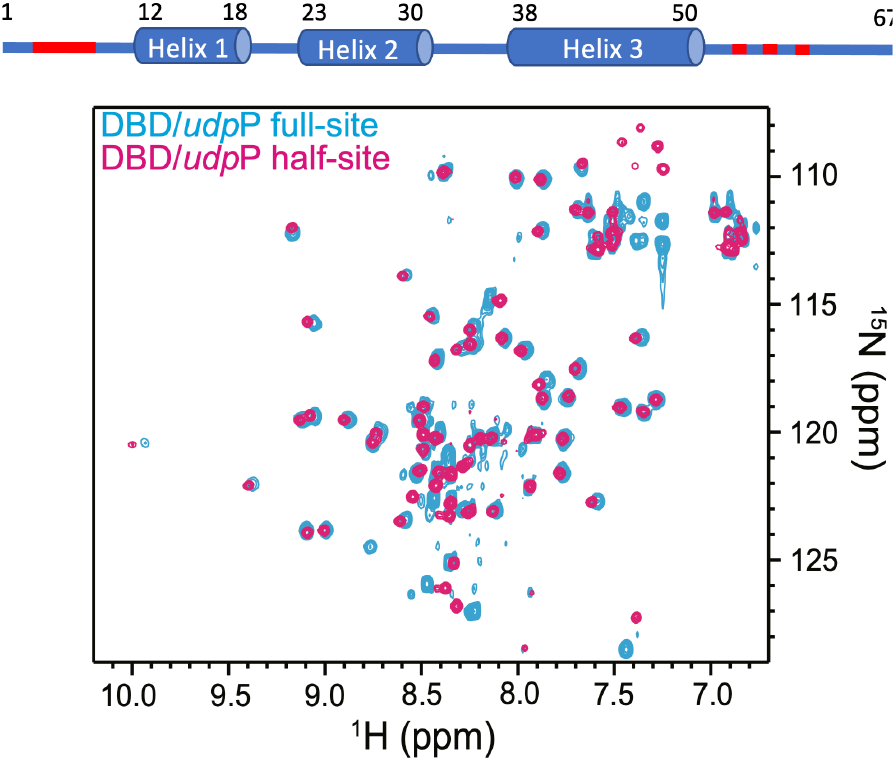
Overlay of ^15^N HSQC spectra for the DBD in the presence of *udp* full-site DNA (cyan) or *udp* half-site DNA (magenta) at 35 °C. The two spectra contain remarkable similarities. The largest changes in peak position are marked red on the sequence diagram. These occur at amino acids 3-8, 54, 58, and 63; these portions at the N- and C-termini were largely found to be unstructured in the DBD/*udp* half-site sample.

**Figure 2.**
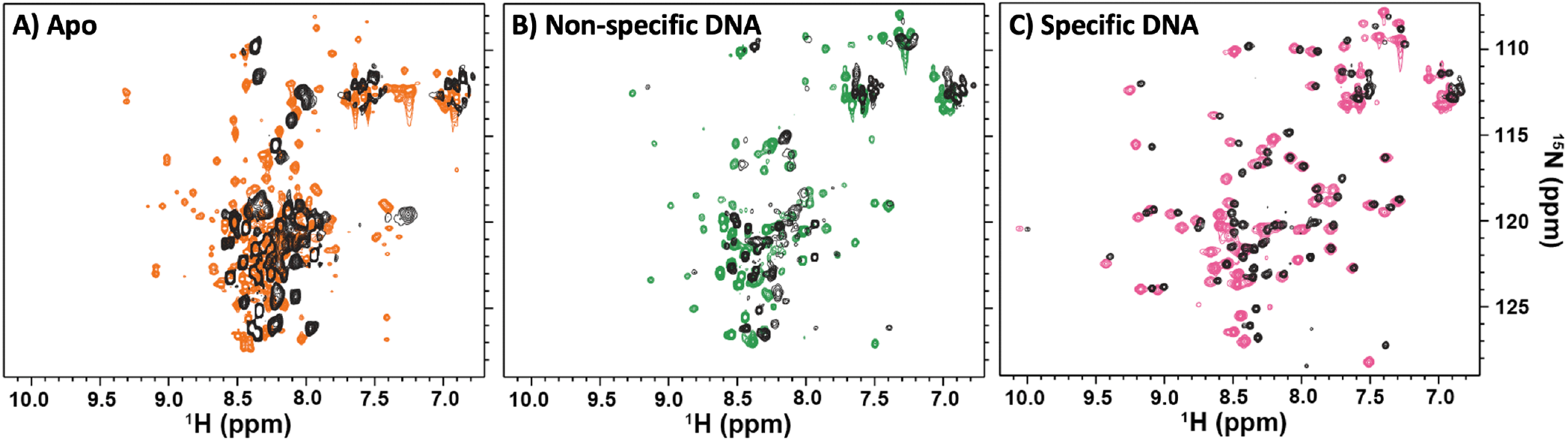
^15^N HSQC spectra of the CytR DBD alone **(A)**, with nonspecific DNA **(B)** and bound to specific DNA **(C)** at two temperatures (colored spectra collected at 20°C; black spectra collected at 35°C). **(A)** the CytR DBD alone exists in multiple conformations at 20°C (orange) and melts to random coil when the temperature is raised to 35°C (black). **(B)** the DBD adopts a single conformation when bound to nonspecific DNA at 20°C (green), but a large part of this structure melts when the temperature is raised to 35°C (black). **(C)** the DBD folds to form a three-helix bundle when bound to specific (*udp* promoter) DNA at 20°C (magenta) and remains folded at 35°C (black).

**Figure 3.**
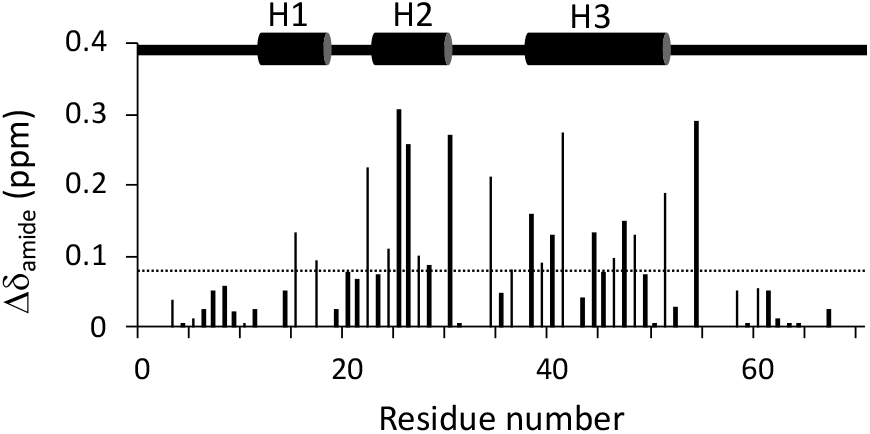
Change in chemical shift (Δδ_amide_) between CytR DBD in the presence of specific DNA (udp half site) and non-specific DNA. The diagram above illustrates secondary structure elements. The dotted line marks a threshold value of 0.09 ppm; changes in chemical shift that can be considered significant. Threshold value was determined by calculating standard deviation of all chemical shift changes.

### Protein/DNA binding affinity measured by fluorescence anisotropy

The spectral differences of the DBD-DNA complexes formed with specific versus nonspecific DNA prompted an analysis of the affinity of DBD for binding to both half-site and full-site operator sequences as well as for binding to two nonspecific DNA oligonucleotides. One nonspecific DNA was the 14 bp oligonucleotide described above; the second was an extended version of this sequence (5’-GTACAGAACATGTCTAAGCATGCTGGGGAC-3’), whose length of 30 bp was chosen to match the *udpP* full-site oligonucleotide. We monitored fluorescence anisotropy of the dye-conjugated DNA substrates to yield an experimental observable for binding. However to analyze the titrations that result, it should be noted that while the fluorescence anisotropy method distinguishes bound from free DNA, it is insensitive to the position in which the DBD is bound to the DNA. It is also relatively insensitive to the stoichiometry and so does not distinguish between a single DBD bound versus multiple DBD’s bound.

**Figure 4.** compares binding of CytR DBD to these four oligonucleotides. The data shown for specific binding to the full-length *udp* operator were reported previously [24]. Analysis according to eq. 3 yields a binding free energy change, ΔG°=-7.14 ± 0.04 kcal/mol, which corresponds to the equilibrium constant for dissociation, k_D_, equal to 4.7 ± 0.3 μM. DBD binding to the *udp* operator left half-site yielded ΔG°=-6.73 ± 0.10 kcal/mol, or k_D_ = 9.6 ± 1.8 μM. Eq. 3 reflects a simple 1:1 binding stoichiometry. The difference of two-fold in apparent affinity between full-site *udp* operator and the left operator half-site is exactly as expected given that the full operator offers two sites for DBD binding. Accounting for the stoichiometry, and assuming no interaction between DBD binding to the left- and right half-sites of the full-length operator, these values are consistent with an intrinsic affinity of 10 μM.

Analysis of the nonspecific binding isotherms using eq. 3 yields ΔG°= −7.70 ± 0.07 kcal/mol (k_D_ = 1.81 μM) for binding to the 30 bp oligonucleotide and ΔG°= −7.03 ± 0.05 kcal/mol (k_D_ = 5.7 μM) for binding to the 14 bp oligonucleotide. The difference in apparent binding free energy ΔΔG°=0.67 kcal/mol, corresponds to a three-fold difference in apparent affinity. Although the site size for nonspecific binding is uncertain, a value of approximately eight base pairs is expected, because this is the length of the inverted repeat sequences for specific binding. With this assumption, the 30 bp oligonucleotide offers three times as many binding sites as the 14 bp oligonucleotide, exactly accounting for the apparent affinity difference observed. Accounting for the stoichiometry these values are consistent with an intrinsic affinity for nonspecific binding (41 μM) that is only four-fold lower than for binding to the *udp* operator. These results indicate that the sequence specificity of the isolated monomeric DBD is modest. The results also show that the DBD is fully bound to both specific and nonspecific DNA under conditions for which NMR spectra are collected.

**Figure 4.**
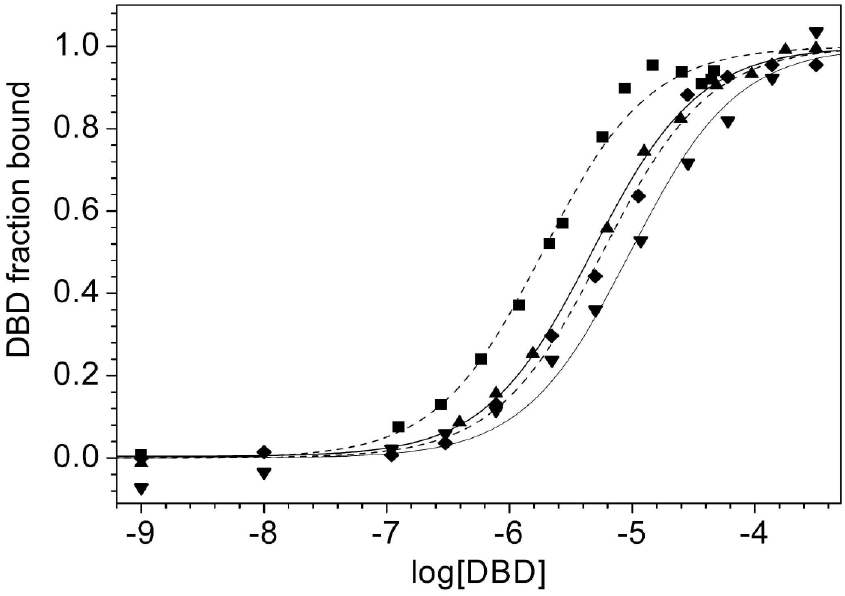
Binding of CytR DBD to udp full-site (filled up triangle) and to its left half-site (filled down triangle) and both 30 bp and 14 bp nonspecific DNA (filled square and filled diamond respectively) at 20 °C, pH 6, as monitored by steady state anisotropy of dye-conjugated oligonucleotides. Solid lines indicate the fitted curves for udp DNAs. Dashed lines indicate the fitted curves for nonspecific DNAs. To facilitate comparison of the binding curves, these have been transformed to a fraction bound scale on the ordinate using the fitted values of r_0_ and r_bound_.

### Structural Analysis of the CytR DBD with Nonspecific DNA

We assessed the structure of the DBD in the presence of nonspecific DNA using NMR spectroscopy. In contrast to both *udp* half and full-site oligonucleotides, many of the ^1^H-^15^N HSQC DBD peak positions in the nonspecific DNA spectrum change dramatically (**Figure 3**). At positions where chemical shifts changes were small or moderate, assignments were extrapolated and confirmed by comparison of sequential ^15^N NOESYHSQC data and deduction based on amino acid content. Analysis of NOE patterns showed strong similarities within the unfolded N- and C-termini of the construct; specifically, amino acids 3-9 and 54-67 (excluding prolines) show both similar NOE patterns regardless of DNA substrate. However, in the intervening regions (10 to 53), most amino acids show very few intermediate- or long-range NOEs in the DBD/nonspecific DNA ^15^N NOESYHSQC spectrum, providing only 36% of the total NOEs compared to the DBD/*udp* half-site (*e*.*g*., **Figure S1**). Notably, the few NOE patterns present in the DBD/nonspecific DNA spectrum do correlate with peaks in the DBD/*udp* half-site spectrum. Unfortunately, these limited NOE data are insufficient to determine the structure of the DBD bound to the nonspecific DNA; however, since we have assigned the backbone amide signals, we can make a quantitative comparison of the dynamics between specific and nonspecific DNA.

### Dynamics of the CytR DBD

Relaxation in the laboratory frame is a useful means to determine motions in the ps-ns timescale range [30]. Having the backbone assignments for DBD bound both specifically and nonspecifically allows us to compare protein dynamics for these two types of interaction. We measured T1 and T2 ^1^H-^15^N relaxation times, and ^1^H-^15^N NOE intensities using 15N HSQC methods. Relaxation and NOE values for the N- and C-terminal ends of the DBD are similar regardless of the DNA binding partner. However, differences occur in the central helical regions when the DBDs interacts with specific (**Figure S2A, C and E**) versus nonspecific (**Figure S2B, D and F**) DNA sequences. **Figure 5** compares the ^1^H-^15^N NOEs when the DBD is bound nonspecifically vs binding to a specific site. Since the ^1^H-^15^N NOE intensity is a sensitive indicator of dynamics, these data show where the DBD becomes more flexible (lower NOE value) when bound to nonspecific DNA. Interestingly, changes in dynamics are not restricted to the loops. The start of helix 2 and middle of helix 3 show substantial changes in NOE intensities that indicate they become more flexible upon binding to nonspecific DNA.

**Figure 5.**
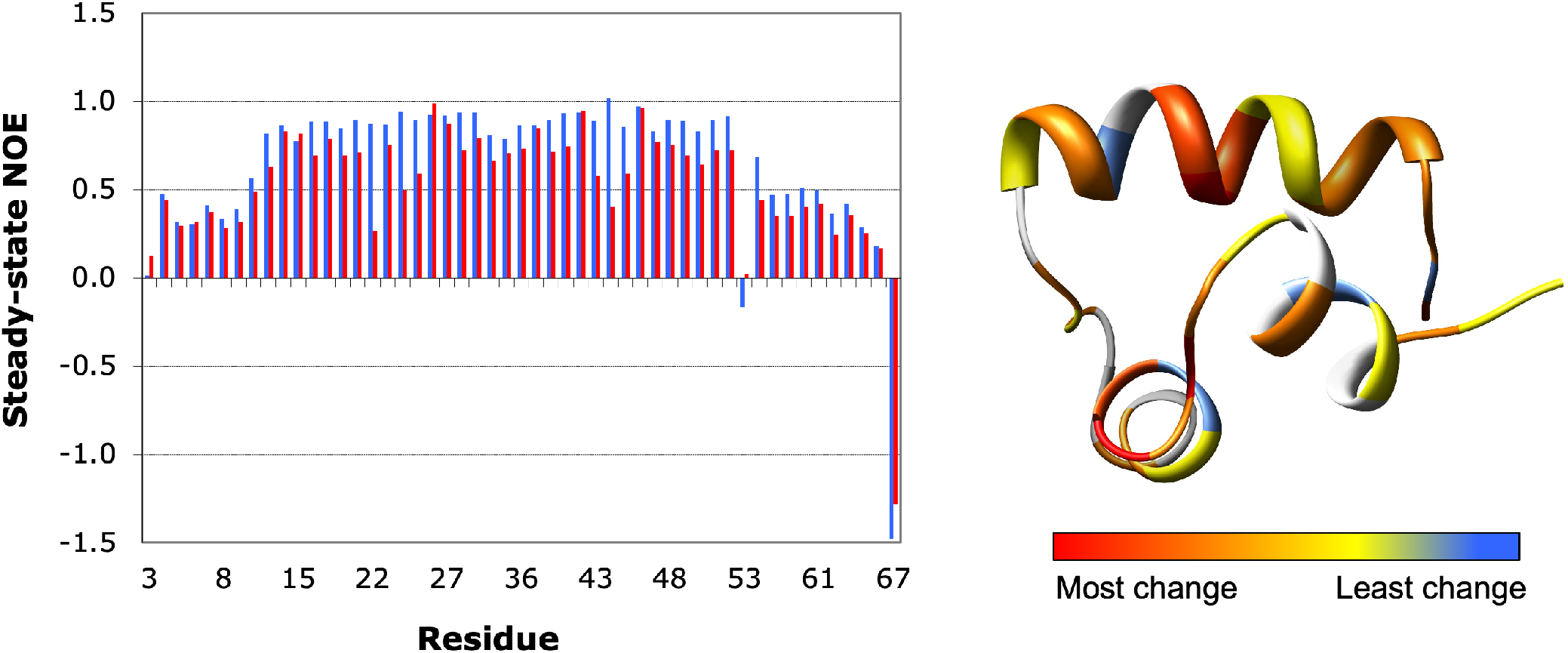
Comparison of steady-state 1H-15N NOEs between the CytR DBD bound to a specific DNA site (blue) and bound to nonspecific DNA (red). Differences in these intensities are mapped onto the DBD structure (residues 9-. The most significant increases in flexibility are at the start of helix 2 and inside helix 3.

**Figure S2B** shows that the DBD T2 relaxation is too rapid to measure using the ^15^N HSQC technique for most of the protein associate with nonspecific DNA. This is consistent with conformational exchange on the NMR timescale for this species. T1 and T2 values on their own are difficult to interpret, but the ratio and product of R2 and R1 (**Figure 6**) provide insight into deviations from the global correlation time. Relaxation rate ratios (R2/R1= [1/T2]/[1/T1]) provide a useful qualitative analysis of dynamics as the average R2/R1 value for a backbone amide in ordered parts of the protein provides a measure of the overall correlation time [31]. R2/R1 can also distinguish between fast (ps-ns) and slow (μs-ms) dynamics [32, 33]. Residues with R2/R1 greater than the limit of one standard deviation from the average value for backbone positions indicate a region of slow (μs-ms) motions and those less than the limit of one standard deviation correlate with fast (ps-ns) dynamics. The R2/R1 ratio for the DBD/nonspecific DNA complex shows that helix 3 appears to exhibit slow motions compared to helices 1 and 2. In addition, R2*R1 can be used to distinguish regions of fast exchange [34]. The R2*R1 product shows chemical exchange at the start of helix 2 and helix 3.

**Figure 6.**
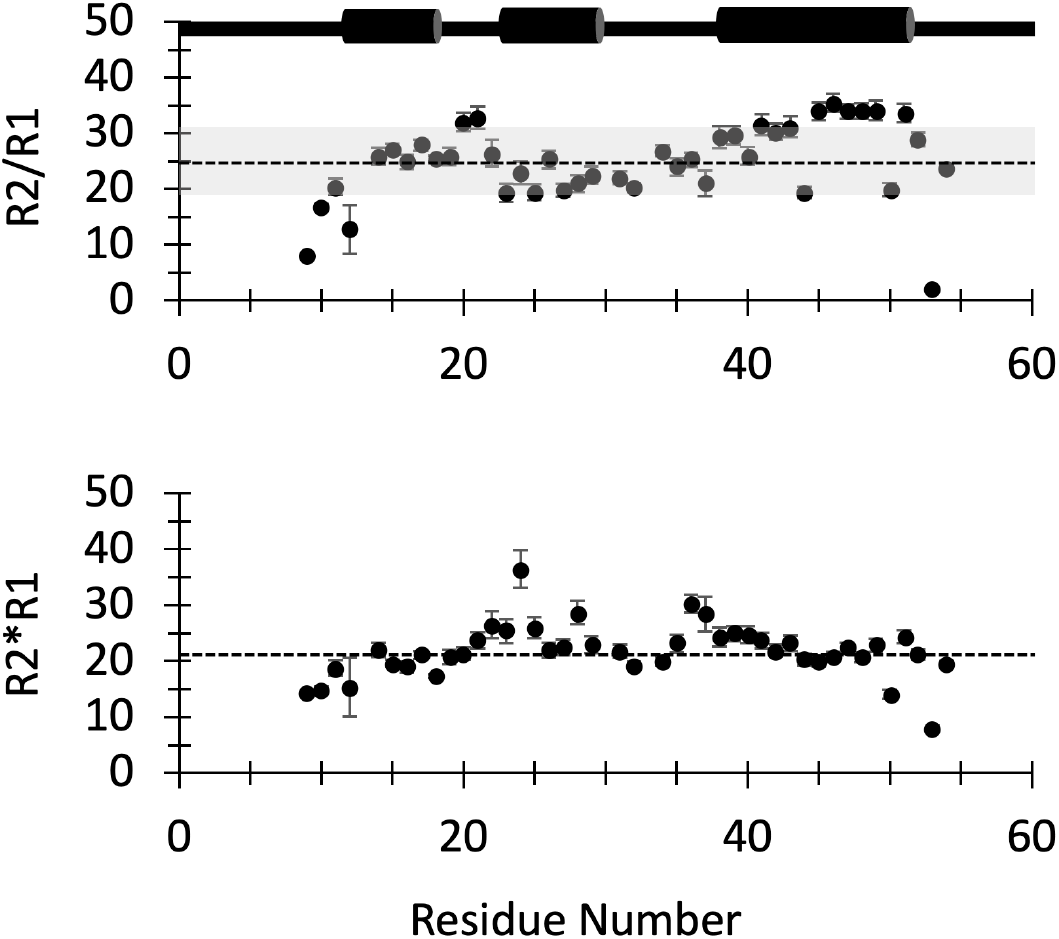
Relaxation rate analysis: R2/R1 and R2*R1 for CytR DBD in the presence of specific DNA (*udp half-site*). The diagram above the graphs corresponds to CytR secondary structure. Dotted lined indicates average values. (A) R2/R1 allows us to distinguish between fast (ps-ns) and slow (μs-ms) motions. (B) R2*R2 highlights chemical exchange above the average; residues 24, 36-37 are the most affected by chemical exchange.

Hydrogen exchange is also a useful measure of protein dynamics and complements relaxation in the laboratory frame by sampling longer timescales of motion. Previously we found complete amide proton exchange occurred within ten minutes of transfer into D_2_O for the free DBD [24]. Notably, we find here that binding to the nonspecific DNA substrate offered no additional protection with complete exchange of the DBD amide protons occurring in the first ten minutes. In contrast, with the DBD bound to the *udp* half-site, we find residues that are protected from exchange for up to three hours following transfer into D_2_O and incubation at 20 °C (pH 6.0 prior to lyophilization). We measured exchange rates and calculated protection factors for all of the positions resistant to exchange. In contrast to the DBD/nonspecific DNA complex that exchanges immediately, all three DBD helices show protection from exchange when complexed with *udp* (**Figure 7A and B**).

**Figure 7.**
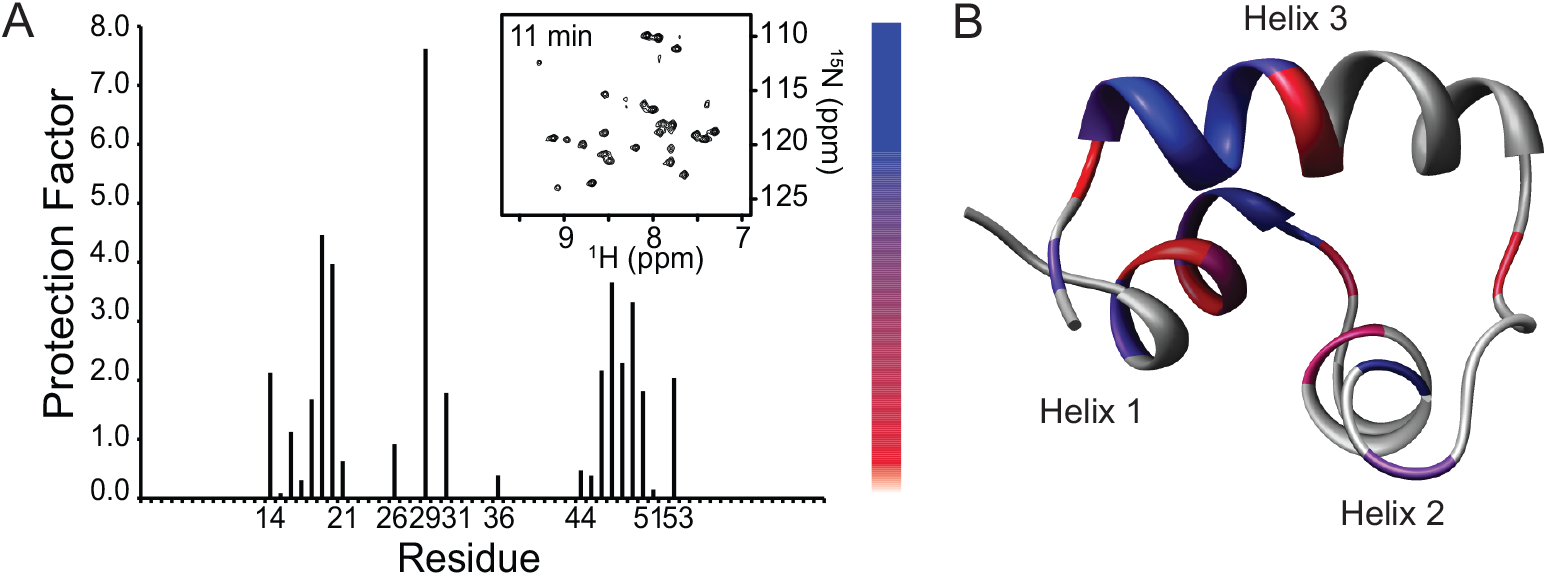
(**A**) Protection factors calculated from hydrogen-deuterium exchange experiments indicate that the first and third helices are most protected from the solvent. Inset HSQC was collected 11 minutes after transfer to D_2_O. (**B**) Hydrogen-deuterium exchange is depicted on a single structure of the ensemble of CytR DBD structures in the presence of *udp* half-site DNA. Blue to red transition indicates high to low protection factors. Residues for which exchange occurred to fast to calculate protection factors are gray. Protection from exchange is primarily isolated to the interface between the first and third helix.

## DISCUSSION

We have now examined the CytR DBD in four states: alone, in the presence of *udp* full-site DNA, in the presence of *udp* half-site DNA, and in the presence of nonspecific DNA. The thermal melting properties of the CytR DBD/DNA systems are intriguing. Although both specific and nonspecific half-site DNA substrates influence the DBD structure by directing the protein into a distinct fold (one set of peaks), the DNA sequences have very different effects on protein thermal stabilities. Significantly, we find that the protein fold is strongly stabilized when bound to specific DNA, but with nonspecific DNA, the DBD is susceptible to thermal denaturation at much lower temperatures, in fact similar to the free state melting.

Our analysis of the DBD spectral features in the presence of the *udp* full-site indicates this species maintains a helix-turn-helix fold similar to the DBD bound to half-site DNA. We find one set of peaks for DBD/*udp* full-site DNA indicating that the protein is in fast exchange between the two recognition sites and consequently we observe an average bound state. Our analysis of DBD NOEs when bound to *udp* full-site DNA reveals that the hinge helix is not formed. We established binding properties of the CytR DBD to the *udp* half-site, as well as to the *udp* full-site. In addition, we studied both nonspecific half- and full-sites. The binding isotherms of CytR DBD to both half-site and full-site DNA are very similar (**Figure 4**); this is consistent with a lack of interaction between DBD monomers when they bind to *udp* full-site. There are two possible interpretations: a) that only a single monomer binds to both the half- and full-site DNA substrates, and b) that the monomers that bind the individual half-sites of the full-site sample do so in an equal and independent manner. Either interpretation supports the relevance of the monomer half-site complex structure presented here. In consideration of regulatory function, equivalent binding constants to half- and full-site DNA combined with the lack of hinge helix formation in full-site DNA suggests that cytidine binding domain (CBD) is necessary for dimer to form. This is either because only the CBD contributes to the subunit-subunit interface or because the hinge peptides require the bottom surface of the CBD dimer to form. The topological differences of CytR and LacR DBDs indicate that the interactions between the surface of the CytR DBD and the CBD should also be unique.

The CytR DBD has similar affinities for both the specific and nonspecific DNA sequences explored in this study. This is not a general feature of HTH DBDs; *e*.*g*., HoxD9 has a 200-fold difference in k_D_’s favoring specific site for binding [35]. Although it appears from fluorescence measurements that all of these DNA substrates bind the DBD with similar affinities, clearly the NMR data reveal stark differences in the protein structure when bound to specific versus nonspecific DNA. *udp* half-site specific DNA can accommodate only one protein molecule when correctly bound. In contrast, protein molecules can bind in multiple positions on nonspecific DNA; collectively, multiple binding sites on a nonspecific substrate could give the appearance that the protein has a higher affinity than individual binding events. This type of binding-mode difference would be consistent with our finding that the structure is stable on specific DNA and much more flexible with nonspecific interactions. However, even in consideration of a lattice model for nonspecific binding, the binding to specific and nonspecific DNAs appear to be energetically similar. At most, specific binding would only be 4-fold higher (∼1 kcal/mol) in the case of protein occupying all available sites on nonspecific DNA.

The energetic balances of the CytR system are quite interesting. Protein folding of the bound conformation must be unfavorable; if not, the DBD would populate the DNA-bound structure even when free in solution. Therefore, favorable interactions with DNA drive unfavorable protein folding for the CytR DBD. Either the two bound protein conformations (specific/nonspecific) are nearly isoenergetic and the intrinsic protein-DNA interactions are nearly isoenergetic (specific vs nonspecific) or the protein-DNA and protein folding are different in specific vs nonspecific cases, but such that they nearly compensate for one another. The DBD folds to a perhaps better defined and more rigid, but energetically less favorable structure in order to make better protein-DNA contacts associated with specific binding, whereas nonspecific binding is inherently weaker and consequently drives less protein folding. The net results are two energetically similar states. Regardless of the apparent similarities in binding affinities, the DNA sequence has a profound impact on the structural properties of the DBD in contrast to LacR.

The results of NMR dynamics experiments indicate significant alterations in the behavior of the protein when bound to a particular DNA substrate. ^15^N relaxation analysis indicates that residues throughout the range 12-50 of the DBD are more flexible in the presence of nonspecific DNA compared to specific DNA, potentially corresponding to movements within the secondary structure. Hydrogen-deuterium exchange reveals that the DBD/nonspecific DNA has greater overall flexibility compared to DBD/*udp* half-site. Despite evidence for a single structure of the DBD with nonspecific DNA, the protein takes on a considerably more dynamic state that allows for the solvent accessibility of all the protein amide groups within the timescale measured here. When bound to nonspecific DNA, larger domain motions could account for the lack of NOEs and consequently would predict that the average structure is different depending on the DNA substrate. In the case of specific DNA, the amide protons of the DBD that were most resistant to exchange are highlighted on the structure (**Figure 7B**).

Although the structure of the CytR DBD resembles the LacR fold in the presence of a specific promoter target, there are differences in the structures induced by nonspecific DNA and the protein free in solution. In the case of LacR, the DBD adopts the same conformation regardless of state (bound or free) [36]. The positions of the first three helices are maintained in all of the LacR structures. Moreover, nonspecific binding does not significantly impact the conformation of the LacR DBD. The PurR DBD structure varies slightly between bound and unbound forms, however structures of the bound form also contain ligand binding domain [37, 38]. CytR is quite different from these structures, adopting a more varied set of conformations depending on the state and binding partner. Although the LacR DBD provided good NOE data to calculate the structure when bound nonspecifically [36], CytR with nonspecific DNA only provides a fraction (1/3) of the NOE constraints needed to define a high-resolution structure. However, having peak assignments allows us to define dynamics on a per-residue basis using NMR techniques. There are significant deviations in protein dynamics between LacR [39] and CytR DBDs. The result of the LacR H-exchange predicts stronger protection factors throughout the protein with helix 2 being most rigid. In contrast, CytR appears to be generally more flexible than LacR, even when bound to a target DNA site and the distribution is different with helices 1 and 3 show more extensive protection from exchange than helix 2.

The distinct structural properties of the CytR DBD free or bound to different DNAs are independent of the interactions with the CBD as this domain is not present in our sample. To generate any effect on differential gene regulation, the CBD interactions with CRP must be involved. Different spacing can affect the conformation of the hinge helices and packing between DBD and CBD; this was anticipated to influence coupling. What we show here is that even without these constraints, the DNA sequence dictates the DBD conformation. Hence, some effect on gene regulation might take place at the level of DBD structure. Therefore, a portion of the differential gene regulation of the full CytR in the cell may be attributed to an altered structure and dynamics of the DNA binding domain itself. Thus, intrinsic affinity for a particular DNA sequence may be overwritten by the structural and dynamical consequences of sequence recognition. In this way, the effects of DNA sequence on the structure and dynamics of the protein may be the basis for distinct levels of gene expression.

Analysis of the structure and dynamics of the CytR DBD allows us to further hypothesize about the link between protein flexibility and DNA recognition. The intrinsic disorder of the free CytR DBD has been confirmed and compared to LacR [7, 13-18]. Our characterization of the nonspecifically bound state classify CytR as forming a fuzzy complex with DNA where the bound protein is only partially structured [19-21]. The dynamics of the DBD may play an integral role in influencing the dynamic state of the rest of the protein. Therefore, the flexibility of the DBD could be transmitting information about the DNA sequence to the CBD and consequently directing the strength of interactions between CytR and CRP. The conformation of the DBD is coupled to DNA binding; however, hinge helix folding and dimer formation requires DBD/CBD interactions, consistent with our hypothesis that differences in spacing are accommodated by changes in protein conformation.

